# A memory of recent oxygen experience switches pheromone valence in *C. elegans*

**DOI:** 10.1101/107524

**Authors:** Lorenz A. Fenk, Mario de Bono

## Abstract

Animals adjust their behavioral priorities according to momentary needs and prior experience. We show that *C. elegans* changes how it processes sensory information according to the oxygen environment it experienced recently. *C.elegans* acclimated to 7% O_2_ are aroused by CO_2_ and repelled by pheromones that attract animals acclimated to 21% O_2_. This behavioral plasticity arises from prolonged activity differences in a circuit that continuously signals O_2_ levels. A sustained change in the activity of O_2_ sensing neurons reprograms the properties of their post-synaptic partners, the RMG hub interneurons. RMG is gap-junctionally coupled to the ASK and ADL pheromone sensors that respectively drive pheromone attraction and repulsion. Prior O_2_ experience has opposite effects on the pheromone responsiveness of these neurons. These circuit changes provide a physiological correlate of altered pheromone valence. Our results suggest *C. elegans* stores a memory of recent O_2_ experience in the RMG circuit and illustrate how a circuit is flexibly sculpted to guide behavioral decisions in a context-dependent manner.

## SIGNIFICANCE STATEMENT

Animals use memories of their recent environment to regulate their behavioral priorities. The basis for this cross-modal, experience-dependent plasticity is poorly understood. *C. elegans* feeds on bacteria in rotting fruit. It monitors O_2_ levels, and switches behavioral state when O_2_ approaches 21%. We show that *C. elegans*’ memory of its recent O_2_ environment reconfigures how it processes sensory information. Pheromones that attract animals acclimated to 21% O_2_ repel animals acclimated to 7% O_2_. O_2_ memory is encoded in the activity history of a circuit that continuously signals O_2_ levels. This circuit is connected to neurons driving pheromone attraction and repulsion. O_2_ experience changes the pheromone responsiveness of these sensors and their post-synaptic targets, correlating with the switch in pheromone valence.

## INTRODUCTION

The body comprises multiple highly integrated subsystems working together to sustain life from moment-to-moment and over long time scales (1). Much of this coordination involves dynamically interacting neural circuits that optimize responses to current circumstances by taking into account sensory input, organismal state, and previous experience (2–8). Circuit crosstalk enables animals to adjust their behavioral priorities in response to a changing environment e.g. variation in temperature, humidity, day length, or oxygen (O_2_) levels (9–13). While some behavioral adjustments can be rapid (14, 15), others develop over time, as animals adapt to changed conditions. How animals store information about their recent environment, and use this information to modify behavioral choices is poorly understood.

The compact nervous system of *Caenorhabditis elegans,* which comprises only 302 uniquely identifiable neurons (wormwiring.org) (16), provides an opportunity to study the links between prior environmental experience, circuit plasticity, and behavioral change. This nematode is adapted to a life feeding on bacteria in rotting fruit (17, 18). It has sensory receptors for odors, tastants, pheromones, and respiratory gases, as well as temperature, mechanical, and noxious cues (19–22). Despite this simplicity, the mechanisms by which its nervous system marshals information about past and present sensory experience to shape behavioral priorities have largely not been dissected. While valuable (23), the anatomical connectome is insufficient to explain or predict neuronal network function (24, 25), partly because neuromodulators can dynamically reconfigure and specify functional circuits (26–28).

When ambient O_2_ approaches 21% *C.elegans* wild isolates become persistently aroused and burrow to escape the surface (29–31). This state switch is driven by tonically signalling O_2_ receptors called URX, AQR and PQR (32, 33) whose activity increases sharply when O_2_ approaches 21% (29, 31, 34, 35). The URX neurons are connected by gap junctions and reciprocal synapses to the RMG interneurons, and tonically stimulate RMG to promote escape from 21% O_2_ (wormwiring.org)(16, 36). URX and RMG are both peptidergic, and at 21% O_2_ tonically release neuropeptides (29, 36). RMG is connected by gap junctions to several other sensory neurons besides URX, including pheromone sensors (16, 37). Whether information communicated from URX to RMG about the O_2_ environment modulates other sensory responses is unknown.

Here, we show that acclimating *C. elegans* to different O_2_ environments gradually reconfigures its response to sensory cues. Animals acclimated to 7% O_2_ but not 21% O_2_ are aroused by CO_2_. Pheromones that attract animals acclimated at 21% O_2_ repel animals acclimated to 7% O_2_. These changes are driven by experience-dependent remodelling of URX O_2_ sensors, RMG interneurons, and the ASK and ADL pheromone sensors.

## RESULTS

### Acclimation to different O_2_ environments reprograms CO_2_ responses

*C. elegans* escape 21% O_2_, which signals that animals are at the surface, and accumulate at 7% O_2_, which indicates that animals are burrowed (29, 31, 32). We speculated that *C. elegans* gradually change their sensory preferences when shifted between these two environments.

To test our hypothesis, we first examined responses to CO_2_. CO_2_ is aversive to *C. elegans*, and its concentration rises as O_2_ levels fall, due to respiration. Animals escaping 21% O_2_ will thus often encounter high CO_2_, creating conflicting drives that we thought could be ecologically significant. Previous work showed that *C. elegans* immediately suppresses CO_2_ avoidance when O_2_ levels approach 21%, due to increased tonic signalling from URX O_2_ sensors (38–41). We speculated that not only current but also prior O_2_ experience remodels *C. elegans’* CO_2_ responses. To test this, we kept wild isolates from California, France, and Hawaii overnight at 21% or 7% O_2_, and compared their responses to 3% CO_2_ on a thin lawn of bacteria kept at 7% O_2_. After halting briefly, animals acclimated to 7% O_2_ became persistently aroused at 3% CO_2_ (**Fig. S1***A–C*), unlike animals acclimated to 21% O_2_.

To probe this plasticity, we studied the N2 lab strain. Unlike natural isolates, N2 is aroused by 3% CO_2_ regardless of prior O_2_ experience (**Fig. 1***A*). In N2, output from the RMG interneurons, a major relay of the circuit signalling 21% O_2_, is blocked by a hyperactive neuropeptide receptor, NPR-1 215V (36, 37). Natural *C. elegans* isolates have a less active receptor, NPR-1 215F, which does not block RMG output (42). Does this account for altered CO_2_ responses? Disrupting *npr-1* caused N2 animals to behave like natural isolates, and to inhibit CO_2_-evoked arousal when acclimated to 21% O_2_ (**Fig. 1***A*). The effects of acclimating *npr-1* animals to 7% O_2_ developed over 16 hours, and were reversed within 3 hours if animals were transferred to 21% O_2_ (**Fig. S**2*A*, *B*). Selectively expressing NPR-1 215V in RMG interneurons prevented *npr-1* animals from acclimating to 21% O_2_ (**Fig. 1***B*), and disrupting the GCY-35 soluble guanylate cyclase, a molecular O_2_ sensor in URX required for the URX – RMG circuit to signal 21% O_2_ (29, 31, 32, 34, 43) had the same effect (**Fig. 1***C*).

**Figure 1.**
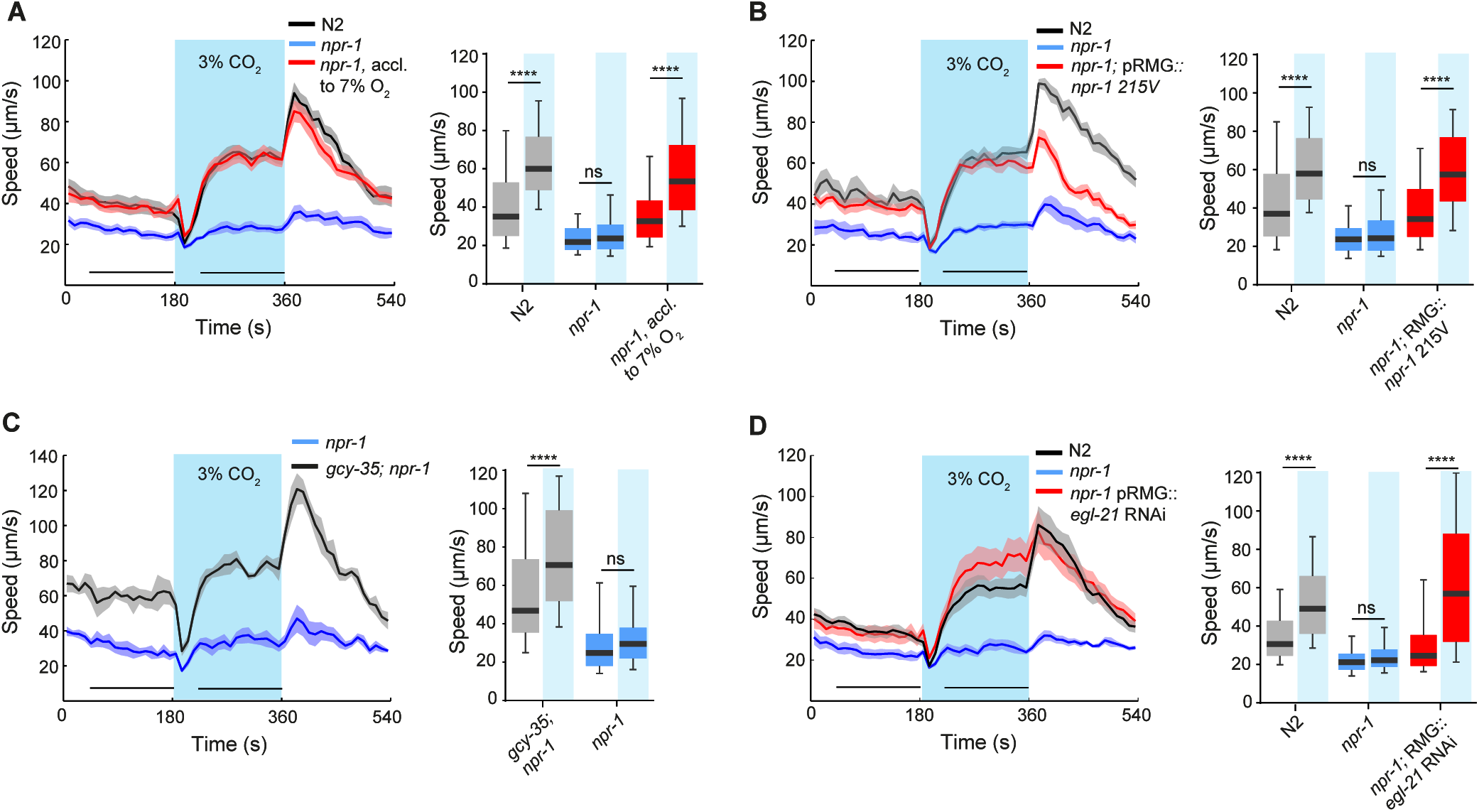
Recent O_2_ experience regulates CO_2_-evoked arousal. (**A**) N2 animals and *npr-1* animals acclimated to 7% O_2_ exhibit a robust and persistent increase in speed when CO_2_ levels rise to 3%, whereas, *npr-1* animals acclimated to 21% O_2_ do not. n = 247–302 animals. **** p < 0.0001; ns, not significant; Wilcoxon signed-rank test. In this and subsequent figures solid lines indicate the mean and shaded areas represent the standard error of the mean (S. E. M). Black bars indicate time intervals used for statistical comparisons (boxplots). Assays were performed in the presence of food and background O_2_ was kept at 7%. (**B**) Selective expression of NPR-1 215V in RMG, (**C**) knocking out *gcy-35*, or (**D**) RNAi-mediated knockdown of the carboxypeptidase E EGL-21 in RMG, prevent *npr-1* animals acclimated to 21% O_2_ from suppressing CO_2_-evoked arousal. n = 104–235. Boxes in this and all subsequent panels show the median (black line) and extend from the 25^th^ to 75^th^ percentiles and whiskers represent 10^th^ to 90^th^ percentiles.

The NPR-1 215V receptor inhibits RMG peptidergic transmission (36). We speculated that circuitry effects of prior O_2_ experience might reflect prolonged differences in RMG peptidergic release. To test this we selectively knocked down the carboxypeptidase E ortholog *egl-21* in RMG using RNAi. Processing of most *C. elegans* neuropeptides depends on EGL-21 (44). RMG-knockdown of *egl-21* prevented *npr-1* animals from acclimating to 21% O_2_ (**Fig. 1***D*). These data suggest that neuropeptide release from RMG is required for *C. elegans* acclimated to 21% O_2_ to suppress CO_2_-evoked arousal.

### Pheromone valence changes with prior O_2_ experience

We studied how prior O_2_ experience alters CO_2_ responses because of the special relationship between these gases. *C. elegans* has, however, many CO_2_-responsive neurons, complicating analysis of how persistent differences in RMG activity alter the CO_2_ circuits (39, 45, 46). Several studies have reported differences in the sensory responses of N2 and *npr-1* mutants associated with altered RMG function (36, 37, 47, 48). We speculated that at least some of these differences could reflect a diminished capacity of N2 animals to acclimate to 21% O_2_ due to reduced neurosecretion from RMG.

One such behavior is pheromone preference (37). Select pheromone blends attract *npr-1* hermaphrodites but repel N2 hermaphrodites (37, 49). We replicated these observations using an equimolar 10 nM mix of asc-ωC3 (ascaroside C3), asc-C6-MK (ascaroside C6), and asc-ΔC9 (ascaroside C9) pheromones, (37, 50–52) (**Fig. 2***A*). We then asked if acclimating N2 and *npr-1* hermaphrodites overnight in different O_2_ environments altered their pheromone response. We assayed animals at 21% O_2_. Whereas *npr-1* animals acclimated to 21% O_2_ were attracted to the pheromone mix, *npr-1* animals acclimated to 7% O_2_ robustly avoided it (**Fig. 2***B*). Acclimating N2 animals at different O_2_ levels did not alter pheromone avoidance (**Fig. 2***B*), recapitulating our observations with CO_2_ (**Fig. 2SC**) Disrupting *gcy-35* switched the pheromone attraction exhibited by *npr-1* animals acclimated at 21% O_2_ into repulsion (**Fig. 2***B*). In summary, reducing the activity of O_2_ sensing circuitry for prolonged periods of time – either via environmental or genetic manipulation – transforms pheromone attraction to pheromone avoidance.

**Figure 2.**
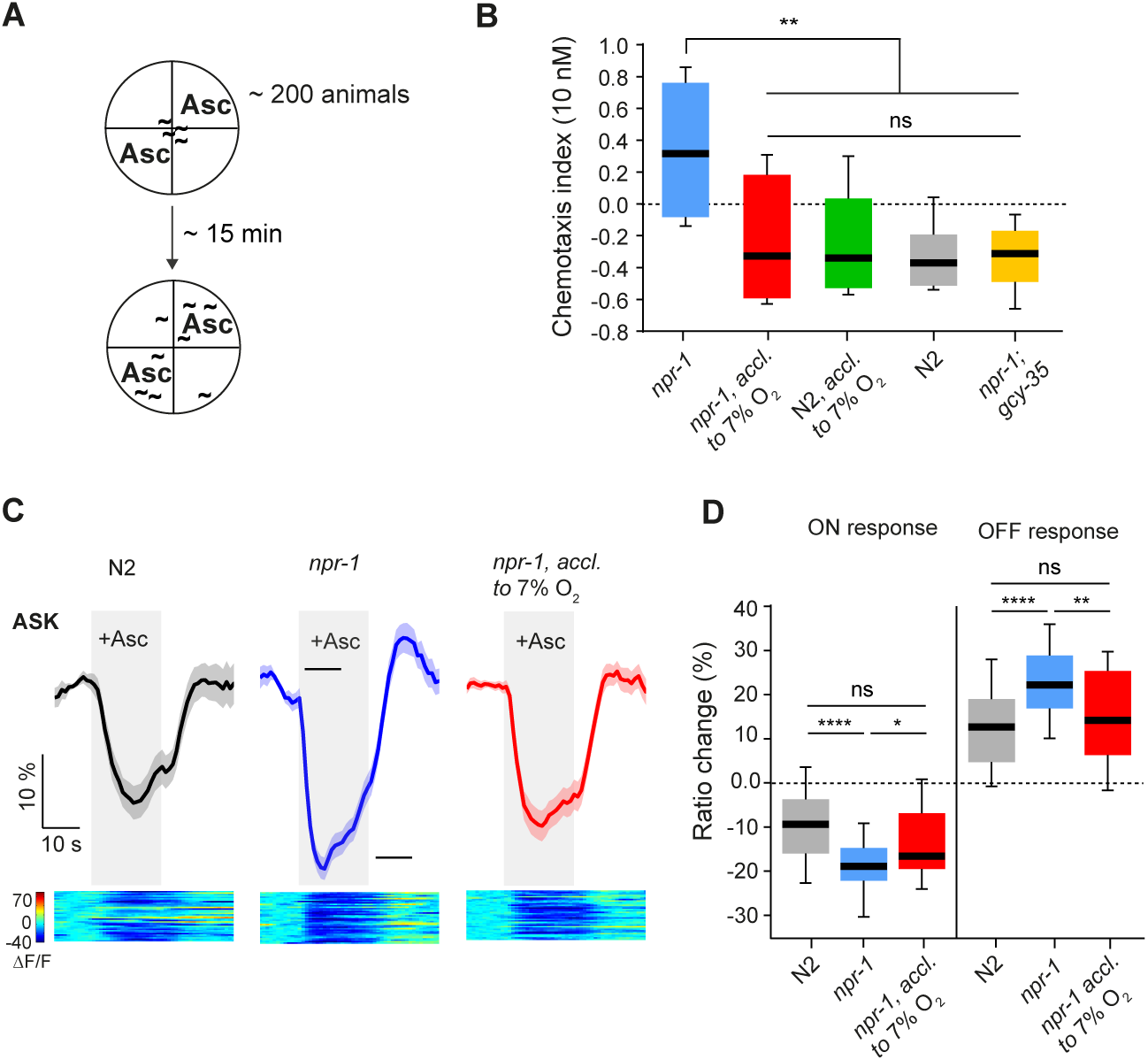
Pheromone valence changes with prior O_2_ experience. (**A**) Quadrant assay for pheromone preference (after (37)). (**B**) Behavioral responses to an equimolar 10 nM mix of C3, C6 and C9 ascaroside pheromones. *npr-1* animals acclimated to 21% O_2_ are attracted to the pheromone whereas siblings acclimated to 7% O_2_ robustly avoid it. N2 avoid pheromones irrespective of whether they have been acclimated to 7% or 21% O_2._ The soluble guanylate cyclase GCY-35 is required for normal O_2_ responses and pheromone attraction in *npr-1* animals acclimated at 21% O_2_. ** p < 0.01; ns, not significant; One-way ANOVA with Tukey’s multiple comparisons test. n = 8 assays each. (**C**) Previous O_2_ experience sculpts pheromone responses in ASK sensory neurons. Acclimation to 7% O_2_ reduces pheromone-evoked Ca^2+^ responses in ASK, consistent with altered behavioral preference. (**D**) Quantification of data shown in (C). Heat maps in this and all subsequent figures show individual Ca^2+^ responses. n = 35–36 animals. **** p < 0.0001; ** p < 0.01; *p < 0.05 ns, not significant; Mann-Whitney *U* test.

### O_2_ experience changes pheromone responses in ASK neurons

How does prior O_2_ experience switch pheromone valence? The altered behavior must reflect some lasting change in the circuitry that couples sensory detection to motor output. The principal neurons driving pheromone attraction are the ASK ciliated head neurons. ASK responds to pheromone with a decrease in Ca^2+^ (the ‘ON’ response) that quickly returns to above baseline when pheromone is removed (the ‘OFF’ response). The pheromone-evoked Ca^2+^ response in ASK is bigger in *npr-1* animals compared to N2 animals, a difference thought to contribute to the opposite pheromone preference of these strains (37). Does prior O_2_ experience change the responsiveness of ASK to pheromones? To test this, we measured pheromone-evoked Ca^2+^ responses in ASK using the ratiometric Ca^2+^ indicator YC3.60. Overnight acclimation at 7% O_2_ attenuated the ASK pheromone response in *npr-1* animals to levels found in N2 (**Fig. 2***C* and 2*D*). Thus, prior O_2_ experience alters ASK pheromone responses, commensurate with a change in behavioural preference.

### Peptidergic feedback heightens RMG responsiveness to 21% O_2_ after sustained exposure to 21% O_2_

RMG interneurons are connected to both the URX O_2_ receptors and the ASK pheromone sensors via gap junctions (wormwiring.org)(16, 37). A simple prediction made by our data is that the response properties of RMG change when *npr-1* animals are acclimated to different O_2_ levels, and this alters the properties of ASK. To explore this we compared the RMG Ca^2+^ responses evoked by 21% O_2_ in *npr-1* animals acclimated to 21% and 7% O_2_. Animals acclimated to 7% O_2_ showed significantly smaller RMG responses than those acclimated to 21% O_2_ (**Fig. 3***A* and 3*B*). Persistent exposure to 21% O_2_ increases RMG responses to this stimulus.

To probe this change in RMG properties we compared the URX Ca^2+^ responses evoked by 21% O_2_ in animals acclimated to 21% and 7% O_2_. URX drives RMG responses (36). URX responses were smaller in animals acclimated to 7% O_2_ (**Fig. 3***C* and 3*D*), suggesting changes in RMG properties partly reflect plasticity in URX. In addition, acclimating *npr-1* animals to 21% O_2_ was unable to increase RMG responsiveness to 21% O_2_ if we selectively knocked down peptidergic transmission from RMG by RNAi of EGL-21 CPE (**Fig. 3***A* and 3*B*). These data suggest there is a positive feedback loop by which tonic peptidergic signalling from RMG in *npr-1* animals kept at 21% O_2_ increases RMG responsiveness to 21% O_2_. Experience-dependent plasticity in RMG and URX represent neural correlates of acclimation to different O_2_ environments.

**Figure 3.**
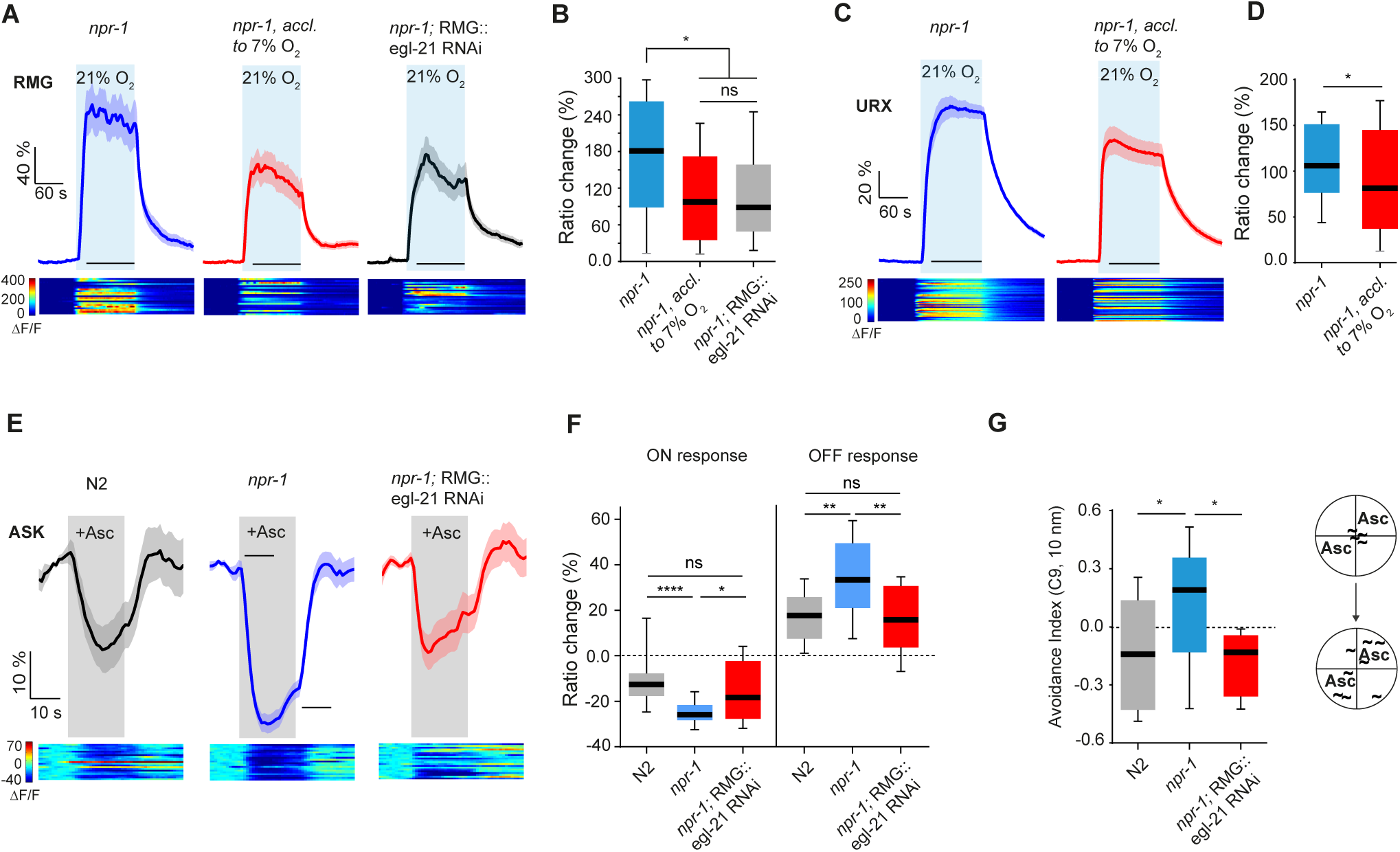
Peptidergic feedback regulates RMG properties and pheromone preference. (**A**) Acclimation to 7% O_2_, or knockdown of *egl-21*, similarly reduce RMG Ca^2+^ responses evoked by a 21% O_2_ stimulus. (**B**) Quantification of data shown in (A). n = 20–21 animals. * p < 0.05; ns, not significant; Mann-Whitney *U* test. (**C**) Acclimation to 7% O_2_ reduces URX Ca^2+^ responses evoked by a 21% O_2_ stimulus. (**D**) Quantification of data shown in (C). n = 38–39 animals. * p < 0.05; Mann-Whitney *U* test. (**E**) Knockdown of *egl-21* in RMG diminishes pheromone-evoked Ca^2+^ responses in ASK to levels observed in N2. (**F**) Quantification of data shown in (E). n = 20–21 animals. * p < 0.05; ** p < 0.01; **** p <0.0001; ns, not significant; Mann-Whitney *U* test. (**G**) RNAi knockdown of *egl-21* in RMG prevents *npr-1* animals acclimated to 21% O_2_ from being attracted to pheromone. n = 12 assays each. * p < 0.05; One-way ANOVA followed by Dunnett’s multiple comparisons test.

Importantly, RNAi knockdown of EGL-21 in RMG altered pheromone responses of *npr-1* animals acclimated to 21% O_2_, reducing pheromone-evoked Ca^2+^ responses in ASK to N2-like levels (**Fig. 3***E* and 3*F*), and conferring robust pheromone avoidance (**Fig. 3***G*). Thus, peptidergic signalling from RMG mediates multiple effects of acclimation to 21% O_2_: an increase tonic Ca^2+^ response to 21% O_2_ in RMG, a bigger ASK response to pheromone cues, and decreased *C. elegans* avoidance of pheromone.

### Communication between neurons in the RMG circuit

The neuroanatomy suggests RMG is gap-junctionally connected to multiple sensory neurons, including ASK, the ADL and ASH nociceptors, the AWB olfactory neurons, and the IL2 chemo/mechanoreceptors (**Fig. 4***A*) (wormwiring.org)(16). Changes in RMG may therefore influence the signalling properties of each of these neurons, and vice-versa. Previous studies suggest that the O_2_-sensing URX neurons cooperate with the nociceptive ADL and ASH neurons, and the ASK pheromone sensors, to promote *C. elegans* aggregation and escape from 21% O_2_ (33, 36, 37, 53). However, in the absence of physiological data it is unclear what information RMG neurons receive or transmit, apart from tonic O_2_ input from URX (29)(36). We asked if O_2_-evoked responses in RMG propagated to ADL and ASK. The wiring diagram suggests ASK and ADL are connected to RMG exclusively via gap junctions. ASK and ADL each showed O_2_-evoked Ca^2+^ responses in *npr-1* animals (**Fig. S3***A*–S3*D*). We also imaged RMG responses evoked by the pheromone mix we used to stimulate ASK (**Fig. 4***B*). RMG responded with Ca^2+^ dynamics similar to those observed in ASK (**Fig. 4***B*), suggesting information can flow from ASK to RMG. These results support a hub-and-spoke model in which different sensory inputs are integrated through gap junctions with the RMG hub (37).

**Figure 4.**
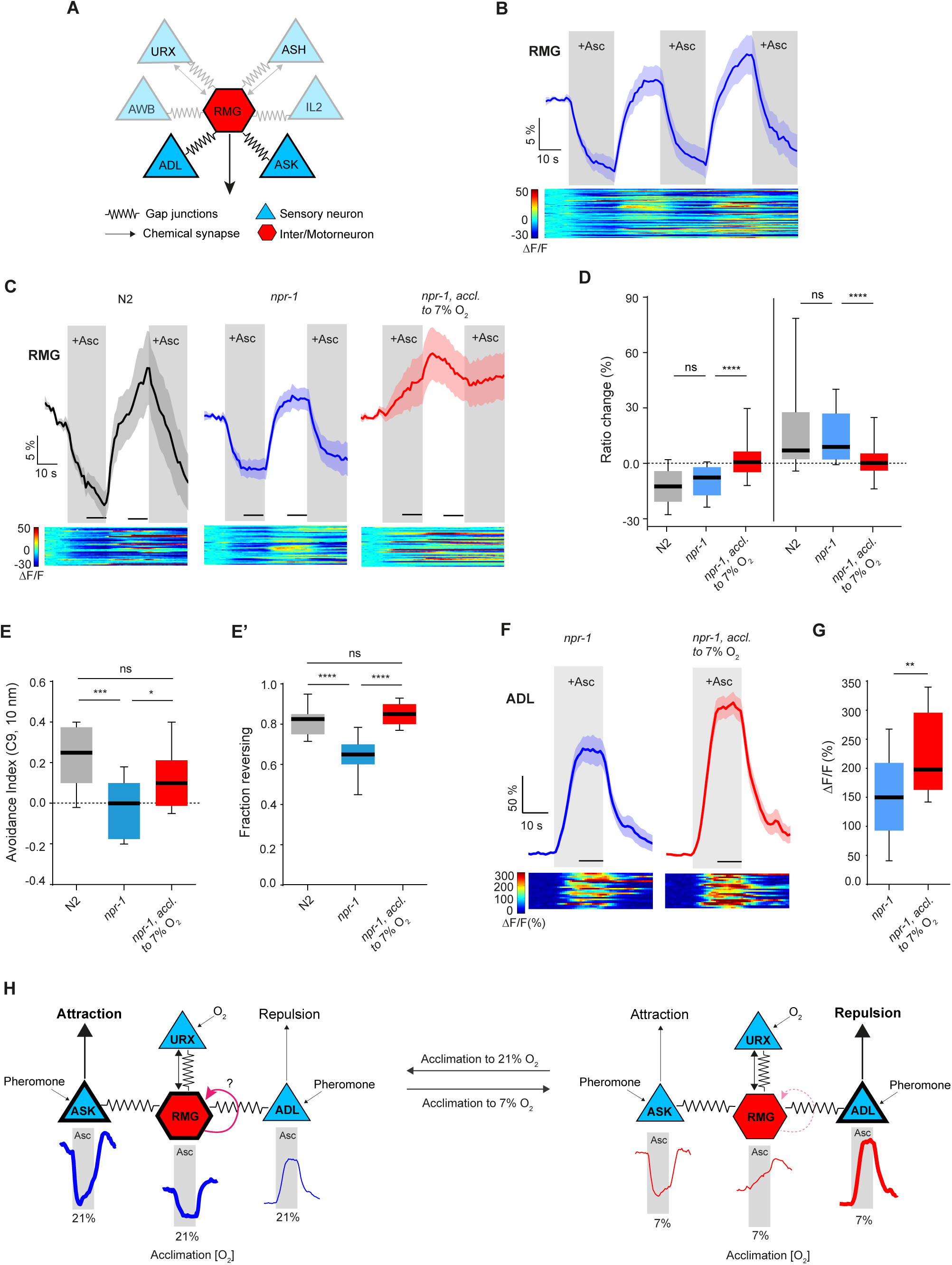
RMG hub neurons respond to pheromones and alter their response according to recent O_2_ experience. (**A**) Circuit showing connections between RMG interneurons and O_2_-sensing, nociceptive, and pheromone-sensing neurons. (**B**) An equimolar (100 nM) mix of C3, C6, and C9 ascarosides inhibit RMG. n = 57 animals. (**C**) RMG shows robust pheromone responses in both *npr-1* and N2 animals. (**D**) Quantification of data shown in (C). n = 35–36 animals. ns, not significant; Mann-Whitney *U* test. (**C**) Acclimation to 7% O_2_ alters RMG properties and diminishes both ON-and OFF-responses to pheromone addition and removal. (**D**) Quantification of data shown in (C). n = 36 animals each. *** p < 0.001; **** p < 0.0001; Mann-Whitney *U* test. (**E–G**) Acclimation to 7% O_2_ enhances ADL pheromone responses and acute pheromone repulsion. *npr-1* animals show decreased avoidance of the C9 ascaroside compared to N2 when grown under standard conditions, but not when acclimated to 7% O_2_. Plotted are the avoidance index (**E**) and fraction of animals reversing (**E’**), in response to a drop of diluted C9 (10 nM) applied to the nose. n = 260–280 animals each. * p < 0.05; *** p < 0.001; **** p < 0.0001; ns, not significant; One-way ANOVA followed by Tukey’s multiple comparisons test. (**F**) The Ca^2+^ responses evoked in ADL by 10 nM C9 pheromone are larger in *npr-1* animals acclimated to 7% O_2_ compared to siblings acclimated at 21% O_2_. (**G**) Quantification of data plotted in (F). n = 23– 24 animals. ** p < 0.01; Mann-Whitney *U* test. (**H**) Model.

NPR-1 215V signalling has been proposed to silence the hub-and-spoke circuit (24, 27). One attractive model is that signalling from the neuropeptide receptor closes RMG gap junctions (37, 49). To investigate this, we first compared pheromone-evoked Ca^2+^ responses in RMG in N2 and *npr-1* animals, but did not observe any significant differences (**Fig. 4***C* and 4*D*). We then compared O_2_-evoked responses in ASK, and also did not observe differences between the two genotypes (**Fig. S3***A* and S3*B*). By contrast, *npr-1* but not N2 animals displayed a strong O_*2*_-evoked response in ADL neurons, (**Fig. S**3*C* and S3*D*); this response, unlike the ADL pheromone response (see below), did not require the TRPV1 ortholog OCR-2 (**Fig. S**4*A*–S4*D*). Although other interpretations are possible, a simple model to explain our data is that NPR-1 215V signalling in RMG affects different gap junctions differently, inhibiting RMG – ADL communication but having smaller or no effects on the RMG - ASK connection.

### O_2_ experience sculpts RMG and ADL pheromone responses

The pheromone attraction mediated by ASK neurons and promoted by RMG signalling is proposed to antagonize pheromone avoidance driven by the ADL neurons in a push-pull mechanism (49). The relative strength of these arms determines the animal’s response. We found that acclimating *npr-1* animals to 7% O_2_ greatly reduced pheromone-evoked responses in RMG compared to animals kept at 21% O_2_ (**Fig. 4***C* and 4*D*). Thus, acclimation to 7% O_2_ weakens both the ASK (**Fig. 2***C* and 2*D*) and RMG circuit elements that drive attraction to pheromone.

Given the neuroanatomy, and the ability of RMG to influence Ca^2+^ in ADL, changes in pheromone-evoked ASK – RMG responses associated with acclimation to different O_2_ levels might alter pheromone-evoked responses in ADL. ADL neurons are activated by the ascaroside C9, and a drop of C9 increases the probability of animals reversing (49). In this behavioral paradigm the fraction of animals reversing provides a measure of the pheromone’s repulsiveness, and is significantly higher in N2 than *npr-1* animals at low pheromone concentrations (10 nM). Higher concentrations of C9 elicit strong repulsion irrespective of *npr-1* genotype (49). We confirmed that N2 animals showed enhanced repulsion from 10 nM C9 compared to *npr-1* animals (**Fig. 4***E*). We then showed that *npr-1* animals acclimated overnight to 7% O_2_ enhanced their avoidance of C9, and behaved indistinguishably from N2 (**Fig. 4***E*). The avoidance index (A.I.) used in this assay (49, 54) is calculated as [(fraction reversing to pheromone) − (fraction reversing to buffer alone)], and any change in the A.I. could reflect an altered response to the buffer rather than to C9. Consistent with enhanced pheromone avoidance, *npr-1* animals reversed more in response to C9 if they were acclimated to 7% O_2_ (Fig. 4E’).

Pheromone-evoked Ca^2+^ responses in ADL neurons were previously characterized using 100 nM C9, a concentration that elicits strong and comparable repulsion in N2 and *npr-1* animals (49). By using GCaMP6 var500 we could record ADL responses to 10 nM C9, and assess the impact of previous O_2_ experience under conditions similar to those used in behavioral assays. *npr-1* animals acclimated to 7% O_2_ showed significantly bigger ADL Ca^2+^ responses compared to siblings acclimated to 21% O_2_ (**Fig. 4***F* and 4*G*). Together our data suggest that acclimation to 7% O_2_ simultaneously weakens the ASK and RMG circuit elements that drive attraction to pheromone, and strengthens the ADL pheromone response driving repulsion, thereby switching the animal’s behavioral choice.

## DISCUSSION

Unfavorable environments can evoke slow, sustained changes in behavioral priorities that reflect an altered internal state. The neural mechanisms mediating such integrative, experience-dependent plasticity are poorly understood. *C. elegans* persistently attempts to escape 21% O_2_ (29), presumably because this O_2_ concentration signals unfavorable surface exposure (31, 32). We find that the O_2_ milieu experienced recently by *C. elegans* changes the way it processes sensory information. Pheromones that attract *C. elegans* acclimated to 21% O_2_ repel animals acclimated to 7% O_2_; 3% CO_2_ triggers sustained arousal in animals acclimated to 7% O_2_ but has comparatively little effect in animals acclimated to 21% O_2_.

A memory of previous O_2_ experience arises from prolonged differences in the activity of a tonically active circuit. Exposure to 21% O_2_ tonically stimulates the URX O_2_ receptor neurons and their synaptic partners, the RMG interneurons. Sustained stimulation of URX and RMG at 21% O_2_ increases their response to 21% O_2_. The reprogramming of RMG requires peptidergic signaling competence in this interneuron. Our data suggest a simple model in which over time, sustained peptide release from RMG at 21% O_2_ feeds back to alter RMG properties. In animals kept at 7% O_2_ peptide release from RMG is low, disrupting the feedback. In this neural integrator model hysteresis in the build up and decay of peptide signaling accounts for the time delays as animals acclimate to 7% or 21% O_2_. We previously showed that neuropeptide expression in RMG is positively coupled to neurosecretion from RMG (Laurent et al., 2015), consistent with a positive feedback loop in this interneuron. Tonic circuit activity is common in brains. We speculate that such circuits will often store information about their activity history, and potentially about the animal’s experience, by incorporating peptidergic positive feedback loops.

RMG has neuroanatomical gap junctions not only with URX, but also with the ASK and ADL pheromone sensors (16, 37). This arrangement suggests that information can be integrated across the circuit (Macosko et al., 2009), but physiological data was hitherto absent. We show that ASK and ADL show O_2_-evoked Ca^2+^ responses, and that acclimating animals to different O_2_ levels alters O_2_ and / or pheromone-evoked responses in each of the URX, RMG, ASK and ADL neurons. Inhibiting peptidergic transmission from RMG prevents RMG and ASK neurons from changing their pheromone responsive properties in animals acclimated to 21% O_2_; it also prevents the experience dependent switch in pheromone valence.

Changes in the pheromone-evoked responses of ASK and ADL neurons are consistent with changes in RMG changing communication across the network. For example, in animals acclimated to 21% O_2_ pheromone-evoked responses in ASK could inhibit ADL pheromone responses, whereas in animals acclimated at 7% O_2_ this communication may be less potent. While this is plausible, we cannot exclude that the intrinsic properties of several neurons in the circuit are altered by O_2_ experience.

We see parallels between our observations and a *Drosophila* study showing that repeated presentation of an aversive shadow cue leads to a persistent change in behavioral state that scales with the number and frequency of the presentations (55, 56). Our findings are also reminiscent of ‘latent modulation’ in the feeding network of *Aplysia*, where the history of activation in some circuit elements has a lasting effect on subsequent responses, most likely by changing neuronal excitability through peptidergic modulation (57, 58).

Why should *C. elegans* reconfigure its sensory responses according to prior O_2_ experience? It is tempting to speculate about a behavioral hierarchy (59) that gives priority to escape from 21% O_2_, and that dominates over sensory drives that could hinder escape from the surface. Animals at the surface may gradually suppress their aversion to CO_2_ to facilitate escape to low O_2_/high CO_2_ environments. Once the threat of exposure at the surface recedes, strong aversive responses to CO_2_ again become adaptive. In a boom-and-bust species like *C. elegans* (18), pheromones may be aversive because they predict an unsustainable population density. However, if escaping the surface is more important than accumulating in a crowded environment, attraction towards pheromones may be transiently adaptive because crowded environments predict reduced O_2_. Irrespective of the precise selective advantage(s), our data suggest *C. elegans* can adopt alternate persistent internal states according to the length of time they have been exposed to aversive or preferred O_2_ levels. In these states neural circuits process sensory information differently, changing the animal’s behavioral priorities.

## ACKOWLEDGEMENTS

We thank Rebecca Butcher for ascarosides, the *Caenorhabditis* Genetics Centre for strains, and members of the de Bono and Schafer labs for advice and comments. This work was supported by the European Research Council (AdG 269058) and the Medical Research Council (UK).

## AUTHOR CONTRIBUTIONS

L.F and M.d. B designed experiments, L.F. performed the experiments, L.F. and M.d.B analysed the data and wrote the manuscript.

## MATERIALS AND METHODS

### Strains

Strains were grown and maintained under standard conditions with *E. coli* OP50 as food (60). To cultivate animals in specific O_2_ environments we used a Coy O_2_ control glove box (Coy, Michigan, USA).

### Behavioral Assays

Locomotion assays were performed as described previously (45, 46). 20–25 adult hermaphrodites were picked to NGM plates seeded 16–20 h earlier with 20 µL of *E*. *coli* OP50 grown in 2 xTY. To create a behavioral arena with a defined atmosphere we lowered a 1 cm × 1 cm × 200 µm deep polydimethylsiloxane (PDMS) chamber on top of the worms, with inlets connected to a PHD 2000 Infusion syringe pump (Harvard apparatus). We pumped in defined gas mixtures (BOC, UK) that were humidified using a sintered gas bubbler (SciLabware, UK) at a flow rate of 3.0 mL/min. Movies were recorded at 2 frames/s using a Point Gray Grasshopper camera mounted on a Leica M165FC dissecting microscope, and were analysed using custom-written Matlab software (36) to detect omega turns and calculate instantaneous speed.

Chemotaxis to pheromones was assayed essentially as described previously (37, 61), using four-quadrant Petri plates (Falcon X plate, Becton Dickinson Labware, USA). For each assay 200 worms were picked to a fresh seeded plate for 2–3 hours, washed three times with chemotaxis buffer, and placed at the centre of a 10 cm assay plate with pheromones in alternating quadrants. Animals were scored after ~ 15 min and a chemotaxis index calculated as (number of animals on pheromone quadrants – number of animals on buffer quadrants) / (total number of animals). We used an equimolar (10 nm) mix of the ascarosides C3, C6, and C9. Assays were repeated on at least 4 different days.

Acute C9 avoidance was examined in the presence of food, using the drop test (62) and as described by Jang and colleagues (49, 54). Responses were scored as reversals if animals initiated a backward movement within 4 s after stimulation that was equal or longer than half their body length. The fraction reversing is given by (number of animals that make a long reversal) / (number if total animals tested); the effect size or avoidance index was calculated as (fraction reversing to pheromone) – (fraction reversing to buffer alone).

### Ca^2+^ Imaging

Ca^2+^ imaging was performed as described previously (37, 49, 63), using microfluidic devices (MicroKosmos, Ann Arbor, MI) to immobilize animals and either a 1:1:1 ratio of three ascarosides (C3, C6, C9) or C9 alone, at the concentration indicated (10nm – nm). For O_2_ and CO_2_ experiments, worms were glued to agarose pads (2% in M9 buffer, 1 mM CaCl_2_) using Dermabond tissue adhesive, with the nose and tail immersed in M9 buffer (45, 46). All imaging was performed on an inverted microscope (Axiovert; Zeiss) with a 40× C-Apochromat lens (water immersion, N.A. 1.0), and MetaMorph acquisition software (Molecular Devices). Recordings were at two frames/s with a 100 ms exposure time. Photobleaching was minimized using optical density filter 2.0 or 1.5. For ratiometric imaging experiments we used an excitation filter (Chroma) to restrict illumination to the cyan channel, and a beam splitter (Optical Insights) to separate the cyan and yellow emission light. Animals were pre-exposed to excitation light for ~ 1 min in all experiments. A custom-written Matlab script was used to analyze image stacks(36).

### Molecular Biology and Generation of Transgenic Lines

Expression constructs were made using the MultiSite Gateway Three-Fragment Vector Construct Kit (Life Technologies). Promoters used include: *sra-9* (3 kb; ASK), *sre-1* (4 kb; ADL), *flp-21* and *ncs-1* (RMG). Promoter fragments were amplified from genomic DNA and cloned in the first position of the Gateway system, genes of interest in the second position, and the *unc-54* 3′ UTR or the SL2∷mCherry sequence in the third position. Constructs were injected at 30–55 ng/µL, with a coinjection marker (*unc-122∷RFP* or *unc-122∷GFP*) at 50–60 ng/µL.

### Statistical methods

Statistical analyses used Prism 6 (GraphPad) and MATLAB (MathWorks). No statistical method was used to predetermine sample size, which are similar to those generally employed in the field. Exact tests used are indicated in figure legends; imaging and locomotion data were analyzed using a non-parametric Mann-Whitney *U* or Wilcoxon signed-rank test.

## SUPPLEMENTAL FIGURES

**Figure S1.**
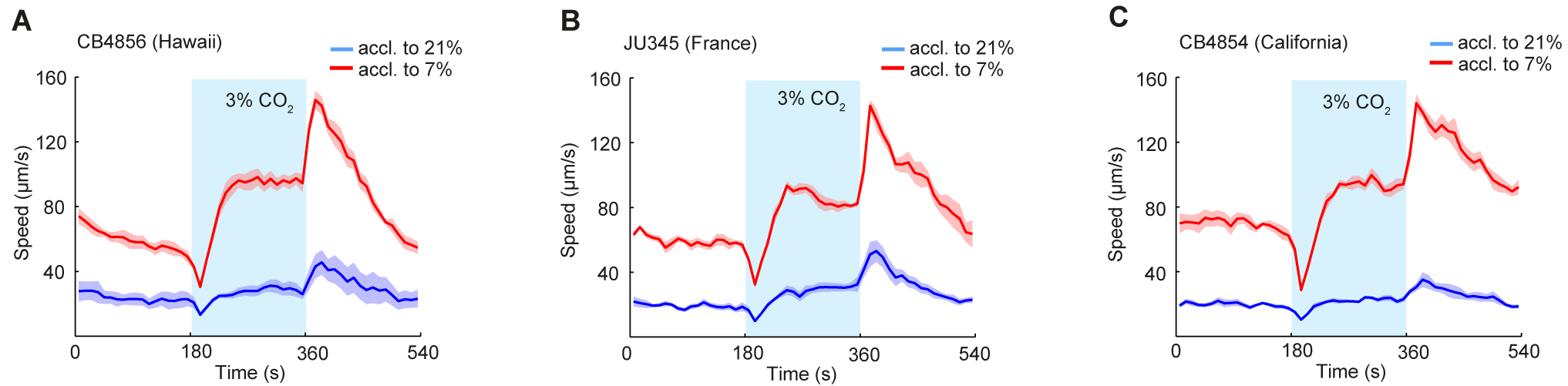
Acclimation to different O_2_ environments reprograms CO_2_ responses in natural *C. elegans* isolates. (**A-C**)Wild strains modulate their CO_2_ response according to recent O_2_ experience. These strains encode NPR-1 215F, the natural low activity isoform of NPR-1. n = 116–171.

**Figure S2.**
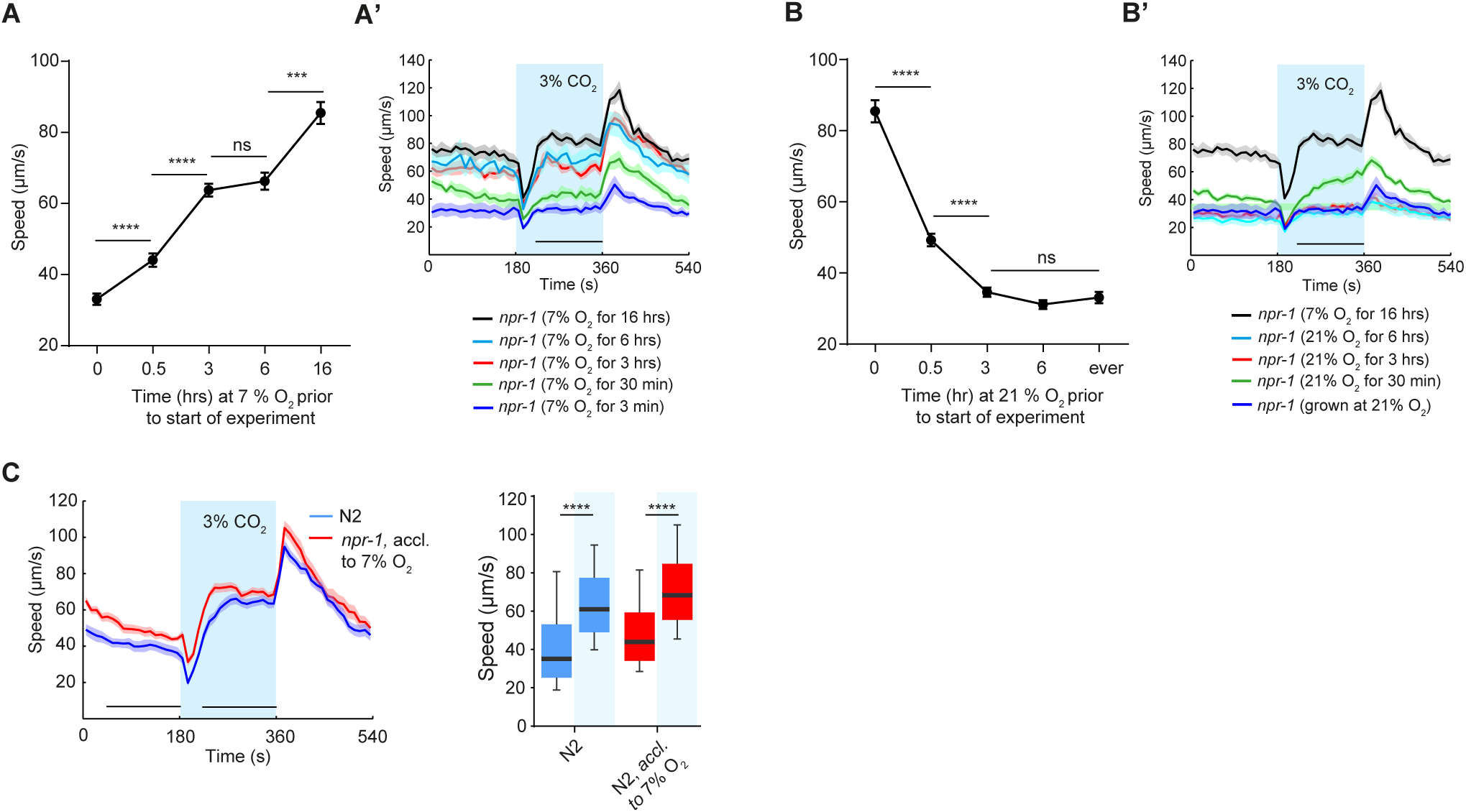
Time line and reversibility of acclimation to different O_2_ levels. (**A** and **B**) Mean speed of *npr-1* animals at 3% CO_2_, plotted against time exposed to 7% or atmospheric (~ 21%) O_2_. (**A**) Acclimation to 7% happens gradually and animals continue to increase their speed over many hours. n = 151–171. (**B**) Acclimation is reversed rapidly, and after ≤ 3 hours animals behave like siblings grown at 21% O_2_. n = 138–188. Error bars in (**A** and **B**) and shaded regions in (**A’** and **B’**) represent S. E. M. **** p < 0.0001; *** p < 0.001; ns, not significant; Kruskal–Wallis ANOVA with Dunn’s multiple comparisons test. (**C**) N2 are strongly aroused by a 3% CO_2_ stimulus, irrespective of whether they have been acclimated at 21% or 7% O_2._ n = 462–518 animals. **** p < 0.0001; Wilcoxon signed-rank test.

**Figure S3.**
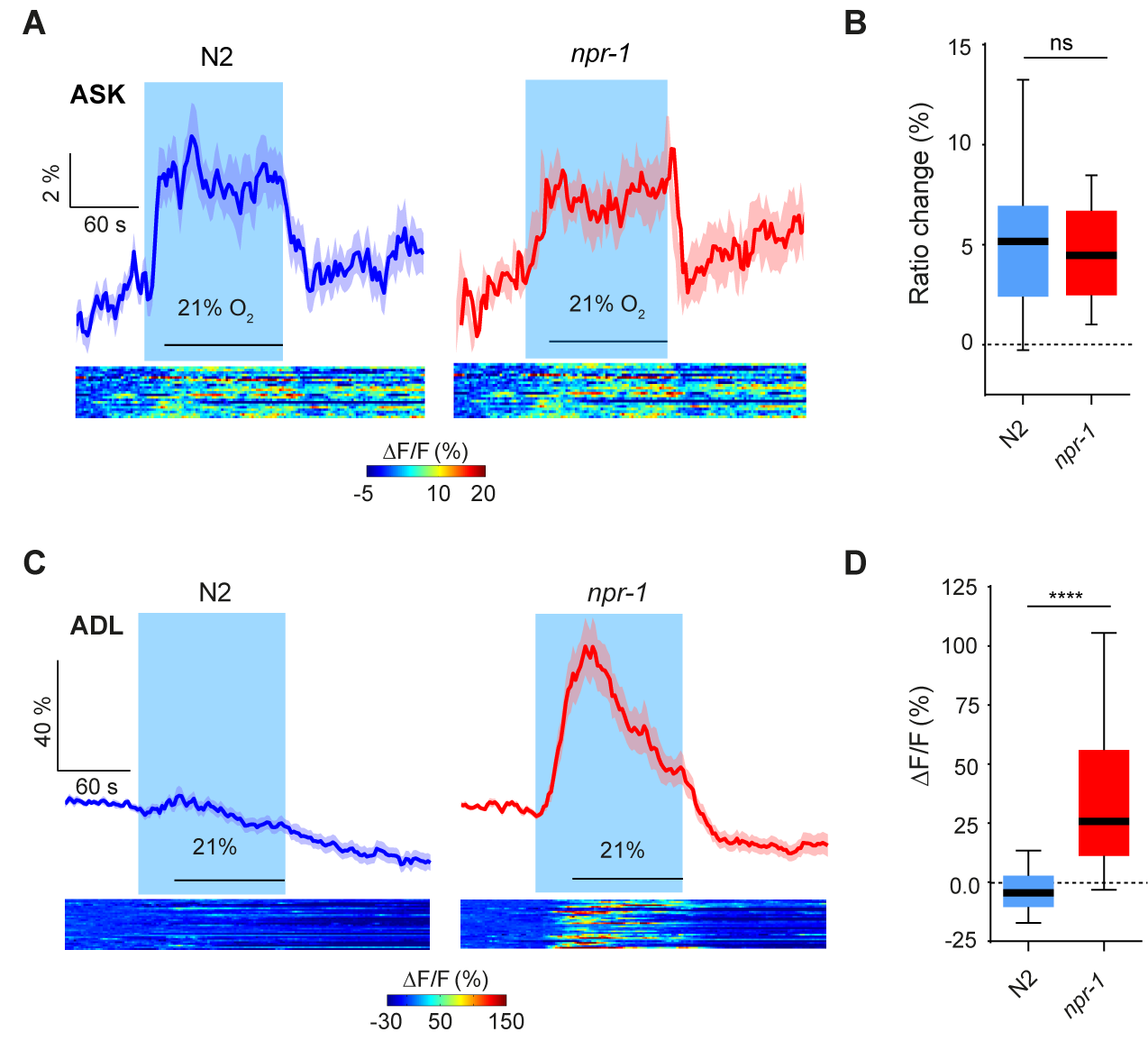
ASK and ADL sensory neuron spokes respond to O_2_. (**A**) O_2_-evoked Ca^2+^ responses in ASK do not differ between N2 and *npr-1* animals. (**B**) Quantification of data plotted in (A). n = 21–24 animals; Mann-Whitney *U* test. Blue shading indicates a shift from 7% to 21% O_2_. (**C**) ADL sensory neurons show robust responses to a 21% O_2_ stimulus in *npr-1* but not in N2 animals. (**D**) Quantification of data shown in (C). n = 30 animals each. **** p < 0.0001; Mann-Whitney *U* test. Note different Ca^2+^ sensors were used to image ASK (YC3.60) and ADL (GCaMP6).

**Figure S4.**
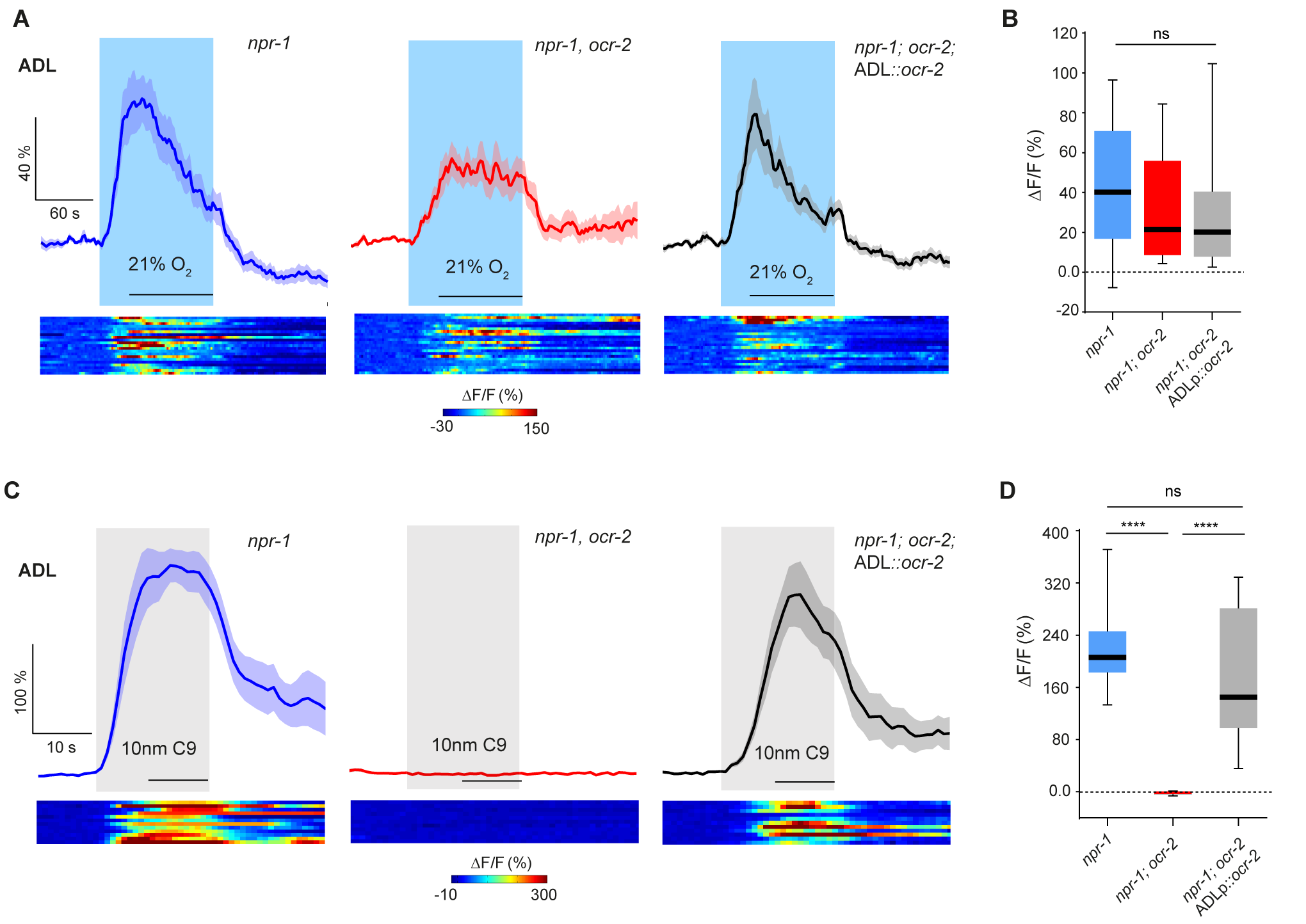
The OCR-2 TRPV channel is required for ADL responses to pheromone C9 but not to 21% O_2_. (**A**) OCR-2 is not required for ADL responses to a 21% O_2_ stimulus, although Ca^2+^ appears to rise less sharply in mutants. (**B**) Quantification of data plotted in (A). n = 21–22 animals each; Mann-Whitney *U* test. Blue shading indicates a shift from 7% to 21% O_2_. (**C**) OCR-2 is required cell autonomously for ADL response to the C9 ascaroside. (**D**) Quantification of data shown in (C). n = 11– 12 animals each. **** p < 0.0001; Mann-Whitney *U* test. Grey shading indicates stimulation with 10 nM C9.

